# RNA 3D modeling with FARFAR2, online

**DOI:** 10.1101/2020.11.26.399451

**Authors:** Andrew Watkins, Rhiju Das

## Abstract

Understanding the three-dimensional structure of an RNA molecule is often essential to understanding its function. Sampling algorithms and energy functions for RNA structure prediction are improving, due to the increasing diversity of structural data available for training statistical potentials and testing structural data, along with a steady supply of blind challenges through the RNA Puzzles initiative. The recent FARFAR2 algorithm enables near-native structure predictions on fairly complex RNA structures, including automated selection of final candidate models and estimation of model accuracy. Here, we describe the use of a publicly available webserver for RNA modeling for realistic scenarios using FARFAR2, available at https://rosie.rosettacommons.org/farfar2. We walk through two cases in some detail: a simple model pseudoknot from the frameshifting element of beet western yellows virus modeled using the “basic interface” to the webserver, and a replication of RNA-Puzzle 20, a metagenomic twister sister ribozyme, using the “advanced interface.” We also describe example runs of FARFAR2 modeling including two kinds of experimental data: a c-di-GMP riboswitch modeled with low resolution restraints from MOHCA-seq experiments and a tandem GA motif modeled with ^1^H NMR chemical shifts.

## 1. Introduction

Noncoding RNA molecules exhibit diverse cellular functions, from catalysis to the detection of small molecules to translation itself [1], and they execute those functions by adopting intricate three-dimensional folds. In such well-defined structures, an RNA’s secondary structure elements are fixed in defined orientations by junctions and tertiary contacts. To keep pace with the acceleration in sequencing technology furnishing new RNA molecules for study, experimental methods for 3D structure determination are being successfully supplemented with structure prediction methods, from physical modeling to knowledge-based techniques [2–5].

Increasingly, new methods for biomolecular modeling are released as webservers, to ensure scientific reproducibility and to mitigate challenges in installation or the availability of computational resources for scientists. This trend includes the ROSIE platform [6, 7], which provides a simplified interface for non-expert users to access computationally intensive protocols developed in the Rosetta framework [8]. The ROSIE server for FARFAR2 enables researchers to model their RNA of interest using a Rosetta algorithm with excellent performance on RNA-Puzzles and other blind challenges [9]. This chapter provides an overview of the two interfaces to this webserver and illustrates how to apply each one to real RNA modeling cases. Because FARFAR2 requires significant computational expense to sample a modeling problem thoroughly, this server, available at https://rosie.rosettacommons.org/farfar2, provides users with a few thousand CPU-hours for their modeling problem, free of charge.

## 2. Method

Here, we illustrate how to use the FARFAR2 ROSIE server in detail using two example problems using the server’s two available interfaces. The “basic” interface allows users to provide nothing more than the two most common piece of data available for RNA structure prediction tasks: the sequence and dot-bracket secondary structure. These data are also the standard inputs provided to RNA 3D modeling webservers from SimRNAweb [10] and RNAComposer [11] to iFoldRNA v2 [12] and MC-FOLD | MC-SYM [13]. No files needs to be prepared. The “advanced” interface permits users to specify significantly more options; every parameter that can affect command-line executions of Rosetta’s FARFAR2 algorithm may be specified through this interface. Users may create an account to receive higher priority and email notifications, or they may submit as guests. Whether or not they create an account, users may make their jobs private if they involve sensitive data.

All files needed to run these examples are available in the Appendix as well as the DasLab/FARFAR2_modeling_examples repository (https://github.com/DasLab/FARFAR2_modeling_examples).

### 2.1 The “basic” interface to the FARFAR2 ROSIE server

First, we examine the basic interface (Figure 1A, B) through interrogation of a pseudoknotted −1 frameshifting element from beet western yellows virus (BWYV; PDB code: 1L2X) [14]

**Figure 1.**
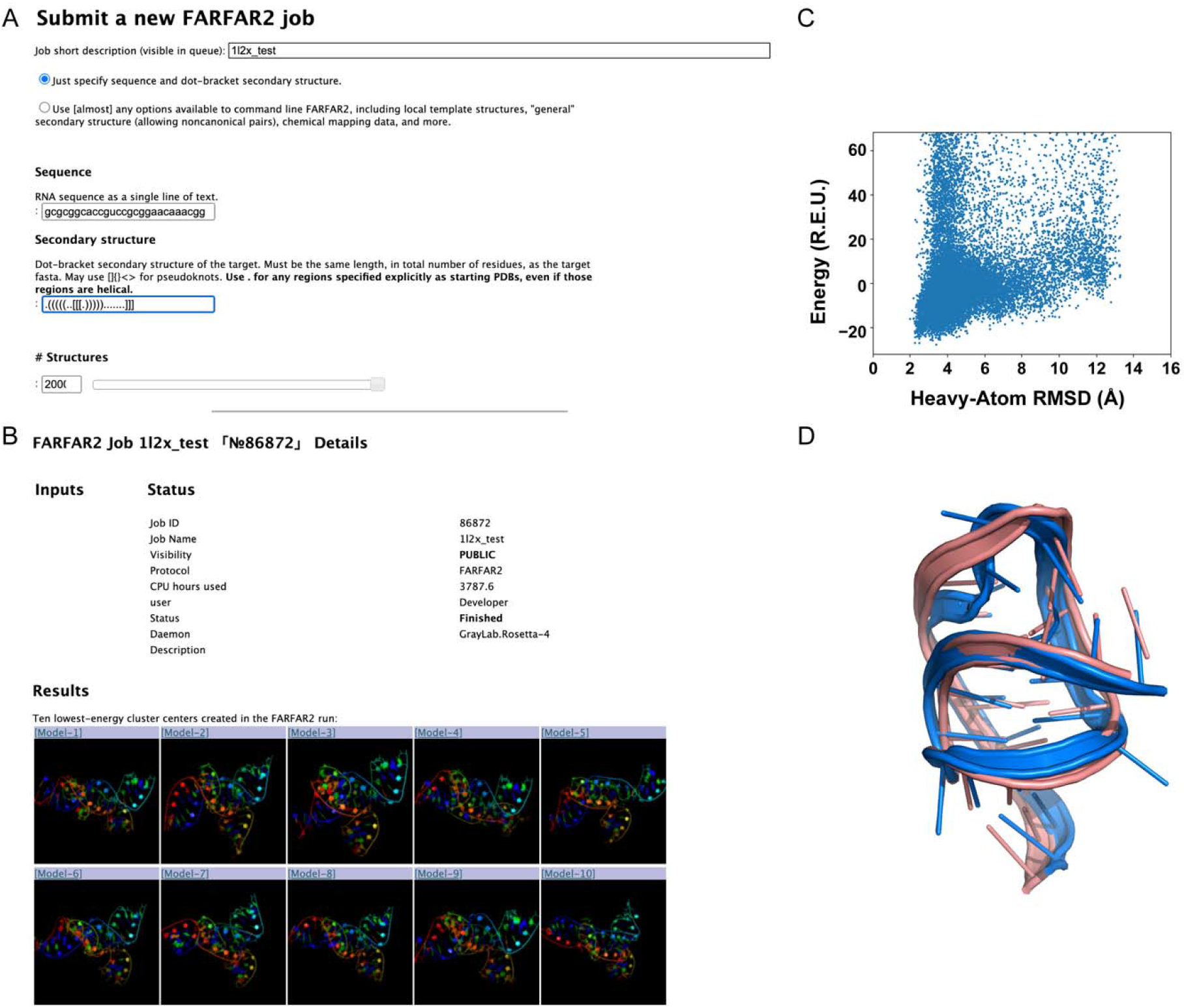
(A) Entering inputs necessary to model the −1 frameshifting element from BWYV to the FARFAR2 ROSIE server’s “basic” interface.”Analysis of 20,000 models resulting from application of FARFAR2 to a viral pseudoknot from beet western yellows virus. (B) Excerpts from the page on the FARFAR2 ROSIE server reporting final results. (C) Plotting the all-atom RMSD to the lowest-energy (in Rosetta Energy Units, or REU) FARFAR2 structure suggests a scoring function that favors a single conformation, and numerous models within 3 Å of this structure. (D) The second-lowest energy cluster (in pink) has an RMSD of only 3.24 Å to the experimental structure (in salmon); in contrast, several other clusters in the top ten are 7-10.5 Å away.

#### 2.1.1. Sequence specification

The user may specify the sequence of the RNA of interest, either as lowercase or uppercase. Chain boundaries ought to be specified via commas. Rosetta’s internal representation of RNA sequence uses lowercase letters, to permit compatibility with uppercase protein sequences in other applications; user specification of capital letters A, C, G, U will be converted to lowercase. For the BWYV frameshifting element segment crystallized and deposited as 1L2X, the sequence input is

~~~
gcgcggcaccguccgcggaacaaacgg
~~~

#### 2.1.2. Secondary structure specification

The user should provide the RNA secondary structure in dot-bracket notation. Rosetta uses a common extension to dot-bracket notation that uses brackets other than parentheses to specify pseudoknots. Pseudoknots through third order may be expressed using matching square [], curly {}, and angled <> brackets. For the frameshifting element of 1L2X, the dot-bracket secondary structure is

~~~
.(((((..[[[.))))).......]]]
~~~

Pseudoknots of higher orders are rare but are found in a handful of structures, such as the 8-stranded nanosquare (PDB code: 3P59) [15]. In these cases, it may be necessary to specify higher order pseudoknots using *matched lowercase letters* from a to z. Critically, because the pairing partners of these letters is ambiguous, each distinct fourth-order pseudoknot must be specified with a distinct letter. As above, any chain boundaries ought to be specified via commas.

#### 2.1.3. Specification of the number of structures generated

There is currently no hard and fast rule for how many structures to generate, but we suggest initial runs start with 1000 models, and increase beyond that if convergence is not achieved (as evaluated in the next section 2.1.4. *Analysis of the resulting structural ensemble)*. The optimal number of models depends on the size and complexity of the modeling problem at hand, the computational resources available, and the RMSD accuracy required for whatever downstream application requires structure modeling. It is possible that there is no way to achieve confident 5.0 Å RMSD predictions on the user’s modeling problem of interest even using a million CPU-hours, and it is possible that their modeling problem is simple enough that the entire space of plausible structures barely spans more than 5.0 Å RMSD (for example, a simple stem-loop). As a baseline heuristic, we generally see significant RMSD convergence over the first 1000 structures generated, even for structures of some complexity, if pseudoknot interactions (as in this example) or other information (tertiary contact templates, experimental data; see below) are available. It is unlikely that FARFAR2 will sample significantly closer-to-native structures past that point. That said, problems as small as the viral pseudoknot RNA in 1L2X are relatively inexpensive in computational cost, so for the purpose of illustrating this example thoroughly, we elect to generate 20,000 structures.

#### 2.1.4. Analysis of the resulting structural ensemble

The FARFAR2 ROSIE server generates several useful analyses from the resulting structures. First, it takes the lowest energy structure from the ensemble, and computes the all-heavy-atom RMSD of each structure to this model. Plotting the resulting ensemble (Figure 1A) helps indicate how cleanly the modeling has converged on a single answer. If there are energetic minima far from the global minimum, this undermines confidence in the modeling and suggests two possibilities. First, sampling may be incomplete, and the true global minimum, yet to be identified, is significantly lower in energy than any minima so far identified. Second, the true minimum may have been identified, but the energy function does not adequately distinguish it from the other minimum structures already identified.

The server also clusters the resulting ensemble and finds the average RMSD among the top 10 cluster centers by energy. This value is known to predict the RMSD to native of the best cluster center by the equation *y* = 0.81 *x* + 3.69 Å, where *y* is the predicted RMSD to native for the closest cluster and *x* is the average pairwise RMSD of the top 10 cluster centers, with an R^2^ of 0.84 [9]. There is a similar relation predicting not just the RMSD error but the uncertainty on this prediction: *yerr* = 0.91 *xerr* + 0.09 Å. In this case, the predicted best RMSD for the BWYV cluster centers is 9.8 ± 1.85 Å. Most of the top 10 clusters indeed have RMSD to the crystal structure 1L2X of 7-10.5 Å, consistent with the predicted accuracy. In this favorable case, the second-best energy cluster turns out to be quite close to the experimental structure, achieving an actual all-heavy-atom RMSD of 3.24 Å (Figure 1B).

### 2.2. The ‘advanced’ interface to the FARFAR2 ROSIE server

Next, we illustrate the use of the advanced interface (Figure 2A, B) on the twister sister ribozyme structure 5Y87 [16], the experimental structure corresponding to RNA-Puzzle 20. During our modeling of this problem for RNA-Puzzles, we made use of a template structure for a hypothesized tertiary contact (a T-loop/intercalation interaction), and the advanced interface is necessary to reproduce this information.

**Figure 2.**
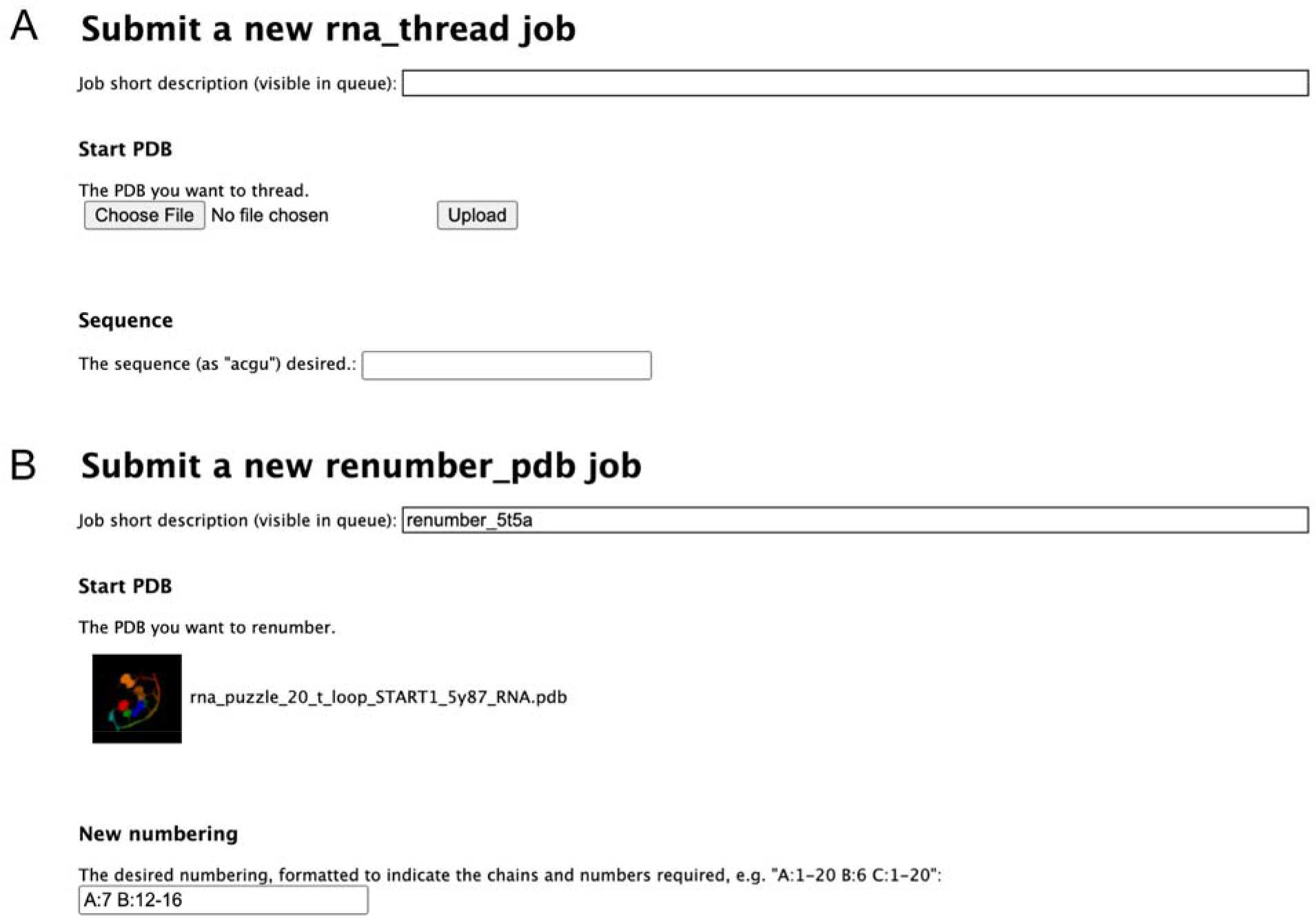
(A) Interface for the *rna_thread* server. (B) Interface for the *renumber_pdb* server, prepared to renumber the starting template as needed.

#### 2.2.1. Sequence specification through a specially formatted FASTA file

We employ a specially formatted FASTA file designed to encode the desired sequence numbering as well as the sequence. The benefits of this FASTA file are significant. First, the specification of custom numbering and chain codes allows the FARFAR2 code to understand desired residue-residue correspondences between models and a provided experimental structure, allowing for the direct computation of native structure RMSD during the simulation. Second, that same correspondence allows for the specification of template structures (see 2.2.6) that give fixed coordinates for particular nucleotides. The FASTA for the 5Y87 modeling challenge is:

~~~
>rna_puzzle_20_t_loop A:1-18
acccgcaaggccgacggc
>rna_puzzle_20_t_loop B:1-50
gccgccgcuggugcaaguccagccacgcuucggcgugggcgcucaugggu
~~~

The FASTA specification allows the study of sequences including chemically modified nucleotides, which must be indicated using special Rosetta nomenclature: indicating the one-letter code as X and specifying a modified base using the PDB 3-letter code, enclosed in brackets. (Thus, the nucleotide dihydrouridine, common in tRNAs, which is found in the PDB as H2U, is indicated in a sequence as `X[H2U]`.) The brackets eliminate potential ambiguity between 3-letter codes and 1-letter codes. The twister sister ribozyme includes no chemically modified nucleotides, so this specific capability is unnecessary.

#### 2.2.2. Secondary structure specification through an uploaded file

In the advanced interface, we supply the secondary structure through a file, rather than a secondary structure string. Unlike when specifying a secondary structure string, the user should not use commas or other characters to separate chains; the FASTA already indicates where chains begin and end.

The secondary structure for our twister sister modeling problem is:

~~~
((((...((((((.(((()))).)(((((.......)))))(((((....)))))))).)
)...))))
~~~

#### 2.2.3. Specification of noncanonical pairs

The ‘ordinary’ secondary structure, as specified above, can contain only Watson-Crick base pairs and G-U wobble pairs. Any base pair indicated above will be assumed to have that standard geometry, and base pairs incapable of a canonical Watson-Crick pairing will prevent job submission. Many noncanonical base pairs nonetheless exhibit highly stereotyped configurations that engage the Hoogsteen or sugar edges of one or both bases, or that engage the Watson-Crick edges in a parallel/trans orientation [17]. For structures known to contain such base pairs, from local motifs like kink turns [18] to tertiary contacts like tetraloop/receptors [19–21], supplementing the secondary structure with this information can be helpful but is rarely available in *de novo* modeling scenarios, so it has not been widely explored with FARFAR2.

Nevertheless, the advanced interface provides two ways to provide noncanonical pair information. First, the user may provide a “general” secondary structure file formatted just the same as above. Any pairs provided in the “general” secondary structure may be satisfied using any combination of nucleobase edges in any orientation, drawn from a database of validated base pairing orientations.

Alternatively, if aspects of the correct noncanonical base pair is known for sure, they may be specified individually in text, formatted like so:

~~~
A:18 A:55 W H A A:24 A:72 X S C
~~~

The string above stipulates that base 18 of chain A and base 55 of chain A must make an antiparallel pair between the Watson-Crick edge of A:18 and the Hoogsteen edge of A:55, and A:24 and A:72 must make a cis pair between any edge of A:24 and the sugar edge of A:72. The permissible base edges are Watson-Crick (W), Hoogsteen (H), sugar (S), and ‘any’ (X), while orientations may be specified as parallel (P)/antiparallel (A) orientation of base normals or through the cis (C)/ trans (T) nomenclature of Leontis and Westhof [17], as well as any (X).

For the RNA-Puzzle 20 twister sister modeling problem 5Y87, there was such a hypothesized set of noncanonical interactions involving a T-loop motif, but this set is actually captured by a local template structure (see step 2.2.6 below), and so specification of noncanonical pairs is unnecessary to reproduce it.

#### 2.2.4. Chain connections

Sometimes there are ambiguities in experimental data intended to guide structure determination. For example, some datasets employing cross-linking or long-distance cleavage information indicate the general proximity of two sets of nucleotides, but no indication as to what nucleotides those could be. “Chain connections” allow users to indicate that there should be a base pair of *some* type – potentially non-canonical – between two sets of nucleotides, without making any assumptions about what the nature or identity of the base pair should be. This is potentially useful for systems with multiple chains, where the “register” of base pairing or tertiary contacts between two chains may be ambiguous. For 5Y87, the secondary structure is known unambiguously thanks to previously published analysis of twister sister ribozymes.

#### 2.2.5. Constraints

Rosetta’s concept of constraints is equivalent to the idea of energetic *restraints* in molecular dynamics. It is common to express certain types of experimental data as energetic restraints that reward a pair of atoms for being a certain distance apart. The FARFAR2 ROSIE server supports the specification of constraint files using Rosetta’s documented constraint file syntax. A specific example related to multiplexed · OH cleavage analysis (MOHCA) experiments is provided in section 2.3.1 below.

The server also accepts two additional parameters governing how constraints are applied. First, users may specify the weight applied to constraints, which permits constraints to influence or outright dominate the energy function. Second, users may alter the way that constraints are applied throughout the course of the low-resolution phase of FARFAR2. Choosing “staged constraints” ensures that constraints between residues that are close to each other in primary sequence are applied earlier in the simulation. Prioritizing local interactions in this way appears to help FARFAR2 discover more solutions that satisfy the constraints.

#### 2.2.6. Input template PDB files

Taking advantage of any known homology of the RNA modeling problem to previously known structural templates accelerates the process significantly and permits a smaller computational expenditure to deliver superior results [22]. The homology does not have to extend over the entire modeled RNA to aid modeling – homology arising from the “modularity” of many RNA motifs, many of which fold to highly similar structures in different contexts [23], is also valuable and illustrated below. FARFAR2 permits the specification of template structures, whose coordinates are kept absolutely fixed during fragment assembly and moved only through energy minimization if desired.

While there can be significant benefits to supplying a template for any junction of an RNA, there are two cases that are especially worth highlighting and have recurred in RNA-puzzles and real-world problems in our lab. First, it can be difficult to model small molecule binding sites with algorithms like FARFAR2 and scoring functions not specialized to that task. But binding sites are often well conserved between RNAs of the same or similar families, and so the junction surrounding, e.g., the S-adenosylmethionine binding site of the SAM-I riboswitch may guide modeling of SAM-IV [9]. Second, tertiary contacts often require precise orientations to form correctly, and during the low-resolution fragment assembly stage the energetic minima are not particularly deep, so even if a tertiary contact is sampled during fragment assembly, only a fraction of models will retain the contact at the end. Thus, local templates of tertiary contacts can enormously focus sampling.

The RNA-Puzzle 20 twister sister modeling problem includes a local template for the intercalated T-loop formed by an adenosine from the catalytic two-way junction and an apical loop distant in the secondary structure. We were able to infer this interaction by a simple analogy to a previously studied twister sister ribozyme, RNA-Puzzle 19, deposited under PDB code 5T5A [24].

We have made additional webservers available for tasks that are essential to manipulating local templates, such as threading on a new sequence (frequently a perfectly valid template will have differences in helix sequence, for example, that do not affect its quality) and renumbering to match the modeling problem at hand. These webservers may be found at https://rosie.rosettacommons.org/rna_thread and https://rosie.rosettacommons.org/renumber_pdb.

To thread a new RNA sequence onto a PDB (Figure 2A), supply the starting PDB and the new desired sequence as ‘acgu’ to the server. The resulting PDB file will have its numbering “reset” to A:1-*N*, where *N* is the total number of nucleotides in your PDB structure. For the twister sister modeling problem, no threading was needed as the sequences were identical in the intercalated T-loop between the template structure 5T5A and the new twister sister RNA.

To update the chains and numbers in a PDB (Figure 2B), supply the starting PDB and the new numbering in the same sort of format used in FASTA files above. That is, chain and residue numbering is described as “A:1” or “B:5” – while multiple consecutive residues with the same chain may be summarized as “A:1-20” or equivalent. We do renumber the T-loop from 5T5A, as it represents an interaction between adenosine residue A:8 and a T-loop comprising A:22-26 in its original context, while in the new twister-sister ribozyme that we seek to model, the interaction is between A:7 and B:12-16, so our input numbering is

~~~
A:7 B:12-16
~~~

The T-loop from 5T5A, renumbered to match the 5Y87 structural context, may be found in the Appendix or in https://github.com/DasLab/FARFAR2_modeling_examples. This template turned out to have 0.36 Å RMSD from the experimental structure.

#### 2.2.7. Alignment PDB structure

There are many situations where the user might have multiple input template structures and an approximate understanding of where they might sit in space. To encode this expectation, the user can supply one “alignment PDB” encoding that understanding. FARFAR2 will impose energetic restraints on each atom in generated models corresponding to an atom in the alignment PDB, penalizing conformations where they stray more than 4.0 Å away from this ideal location. Thus, models will reliably have the desired orientation, but small deviations necessary to accommodate sequence context changes from the template may be permitted.

This feature is also useful for understanding the best possible models achievable by FARFAR2 that are close to a target structure like an experimental structure. Comparison of energies of such near-native models with unrestrained de novo modeling has been useful in understanding limitations in the FARFAR2 energy function [9].

#### 2.2.8. Native PDB structure

The sequence and numbering of any specified native structure must correspond exactly to the provided FASTA file. This structure will of course be unavailable for actual blind challenges approached using the webserver, but for benchmarking cases like the twister sister ribozyme, it is available (5Y87) and we use it here in this example. This file is also supplied in the Appendix as well as the DasLab/FARFAR2_modeling_examples repository (https://github.com/DasLab/FARFAR2_modeling_examples).

#### 2.2.9. High resolution minimization settings

Following fragment assembly, FARFAR2 refines structures through continuous minimization of torsion angles in an all-atom scoring function. This is not strictly mandatory, but structures that have not been optimized in this way will possess nonphysical chain breaks and clashes. If minimization is desired (it is active by default), then users may select one of three high-resolution energy functions. The default is the best-performing setting and the standard for FARFAR2.

By default, residues drawn from template structures are not minimized. Users may additionally specify residues from input template structure that may move during minimization. The input format indicates each residue by residue number and chain, like “`A:1`”; a series of residues may be specified as “`A:1-3 A:5`” and so on. This option is especially useful when the new modeling context for a template is somewhat different from its original context. Because we used an existing twister sister ribozyme structure for the intercalated T-loop template in this modeling, we did not use this option, but if we had used a different structure with a T-loop – say, a tRNA – it may have been helpful.

#### 2.2.10. Low-resolution fragment assembly settings

The initial low-resolution fragment assembly stage also has several manipulable parameters. The user can raise or lower the simulation temperature, which affects how likely a fragment move is to be accepted. They may select the current, updated fragment database; the previous Rosetta default in effect from 2010-2019; or the original fragments that only used one structure of an E. coli 23S rRNA. They may enrich the existing fragment database by adding in torsional combinations drawn from a Gaussian centered at the torsions of each experimental fragment. Additionally, they may enforce an approximate symmetry condition: for example, if the user is interested in modeling a duplex, then they may wish to ensure that any fragment move is applied concurrently to each strand. These non-default options have not been explored widely.

The final condition that modifies the low-resolution phase can be important for rigorous benchmarking: an option originally added to Rosetta when the Das lab needed a standard for comparison in stepwise Monte Carlo benchmarking [25]. The user may choose to *exclude homologous fragments from the fragment library* that resemble the native structure too closely. Using a structure of a twister sister ribozyme like 5Y87 as a benchmark case for testing FARFAR2 would be unfair if fragments from that same ribozyme were present in the library. The fragment exclusion algorithm looks at every 6-mer sequence in the native PDB and removes any fragments that match those 6-mers in sequence and whose substitution would result in a conformation closer than the provided RMSD radius to the native conformation. The sequence match for what might get removed defaults to matching purine/pyrimidine identity but could either ignore sequence entirely or require an exact sequence match. For RNA-Puzzle 20, PDB 5Y87, we supply an RMSD radius of 1.2 Å, following the FARFAR2 study [9].

#### 2.2.11. Experimental data

The user may have external experimental data to guide the modeling process. NMR chemical shifts may be specified using a variant on the STAR 2.1 format. Details of the full CS-ROSETTA-RNA protocol [26], is extensively documented in Rosetta demos available at https://www.rosettacommons.org/demos/latest/public/cs_rosetta_rna/README and application documentation at https://www.rosettacommons.org/docs/latest/application_documentation/rna/CS-Rosetta-RNA, but augmenting FARFAR2 modeling with chemical shift scoring requires only the specification of a STAR 2.1 format chemical shifts file as described here. Section 2.3 gives an example.

#### 2.2.12. Number of structures to generate

The number of structures needed for a whole RNA structure, varies considerably with its size and complexity. The presence of the intercalated T-loop tertiary contact aids the modeling, as that tertiary contact is sufficient to give the whole RNA a globular fold and restricts the possible low-energy structures. In our original blind challenge effort, and in our subsequent simulated benchmark, we were able to generate structures closer than 4.0 Å RMSD to the crystal structure with only a few thousand models. For the RNA Puzzle 20 twister sister ribozyme 5Y87, we generate 20,000 models, simply to provide a thorough exploration of the modeling problem for this work.

#### 2.2.13. Analyze the results

The FARFAR2 ROSIE server (Figure 3A) provides a set of data for this twister sister problem similar to the first pseudoknot benchmark case, albeit with some small differences. Because we provided a native PDB structure in this example, the server does not rescore the ensemble to the lowest energy model. Instead, it plots the score against RMSD to the provided native structure. In part, we can attribute the method’s success on a modeling problem of this complexity to the high similarity in T-loop conformation between the template and target structure (Figure 3B). The resulting ensemble (Figure 3C) may be interpreted similarly to the simpler pseudoknot modeling problem above (Figure 1), albeit with the certainty that the lowest RMSD structures are the most native-like due to the specification of the native structure as a reference here. So energetic minima far from the native structure must indicate scoring function issues. In this case, the resulting top 10 cluster centers automatically generated by the webserver have average inter-model RMSD of 7.6 Å, indicating 9.9 ± 1.9 Å minimum cluster RMSD to native. In fact, the clusters are each quite similar to the experimental structure and range from 5.3 – 9.3 Å, and the best cluster has a significantly superior RMSD, at only 3.91 Å RMSD (Figure 3D). This suggests that the use of such an accurate template significantly helped structure prediction exceed typical expectations.

**Figure 3.**
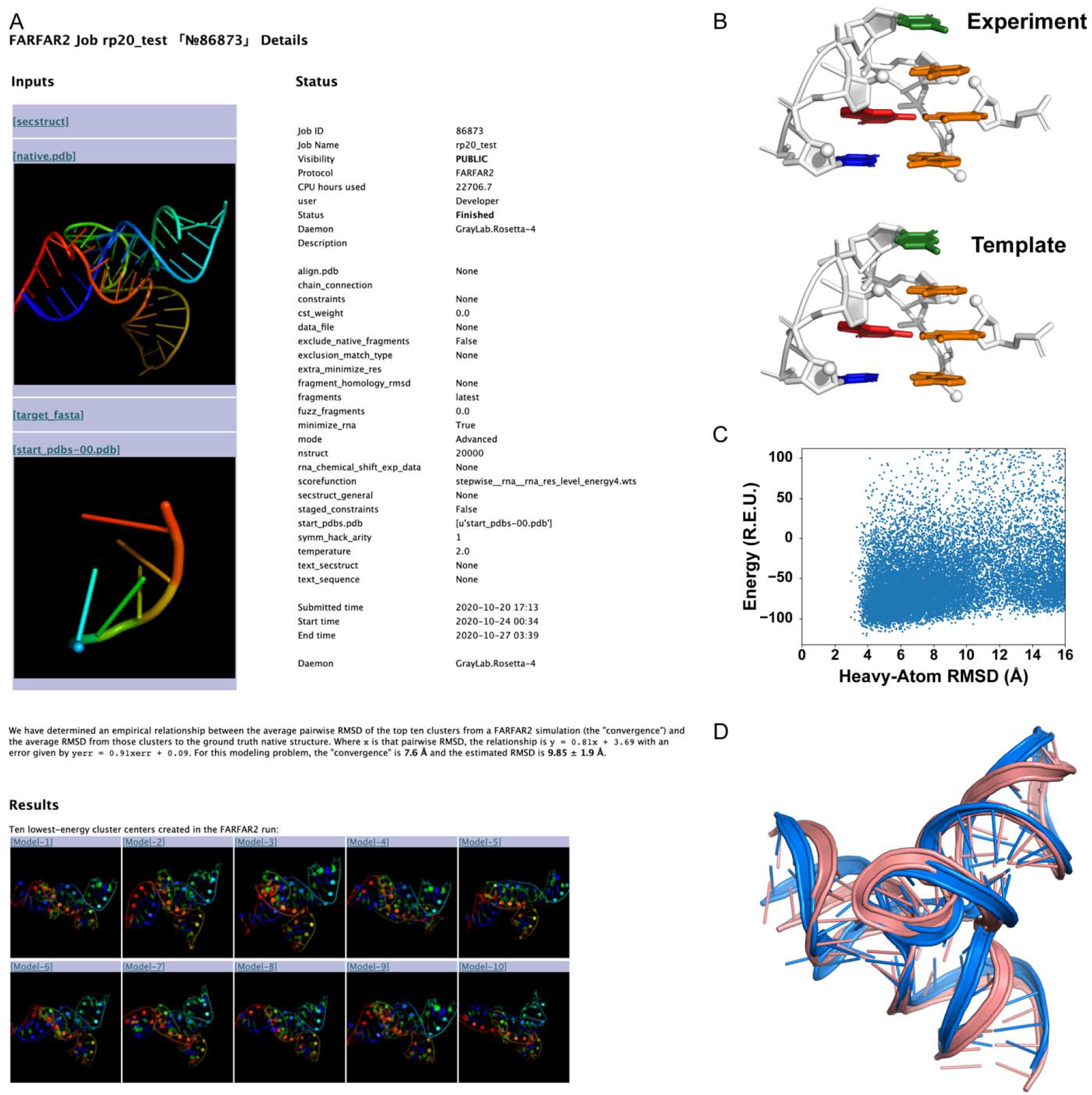
(A) The job submission page that appears upon submitting the suggested inputs to simulate RNA Puzzle 20. (B) A close-up comparison of the excellent template T-loop structure from 5T5A, as compared to the experimental coordinates for the target ribozyme 5Y87. (C) Analysis of the ensemble of FARFAR2 structures generated.

Plotting the all-atom RMSD to the lowest-energy structure suggests good sampling and a scoring function that favors a single conformation. (D) While each cluster center is each fairly close to the experimental conformation (in marine), one of them (in pink) is especially near, with an RMSD of 3.91 Å.

### 2.3. Additional illustrations of advanced interface: experimental data

Experimental data can dramatically improve convergence and accuracy of RNA modeling. The FARFAR2 ROSIE server is well-equipped to handle two kinds of experimental data, multiplexed · OH cleavage analysis (MOHCA) and NMR ^1^H chemical shift data, briefly discussed here. (The recent Ribosolve pipeline integrates Rosetta RNA *de novo* modeling with cryo-EM; a separate ROSIE server is under development for that application and is not described here.)

#### 2.3.1 MOHCA-seq with FARFAR2

MOHCA [27] and MOHCA-seq [28] experiments use tethered hydroxyl radical sources and sequencing readouts to discover “strong” and “weak” signals of nucleotide-nucleotide proximity. These signals can be used to guide FARFAR2 modeling; each signal results in a restraint expressed as the sum of two functions, whose weights are given by the strength of the constraint. A “strong” restraint between chain A nucleotides 2 and 38 at their O2’ and C4’ atoms would be specified via:

~~~
AtomPair O2′ 2A C4′ 38A FADE 0 30 15 −4.00 4.00
AtomPair O2′ 2A C4′ 38A FADE −99 60 30 −36.00 36.00
~~~

Omission of the chain letter leads Rosetta to interpret the sequence position as an absolute number within the PDB (i.e., sequentially starting from 1), which may not match the numbering of the user’s PDB. The parameters for the FADE constraint are described in https://www.rosettacommons.org/docs/latest/rosetta_basics/file_types/constraintfile. The specification above gives a penalty smoothly ramping up to 4 Rosetta energy units as the inter-atom distance shifts away from 15 Å down to 0 Å and up to 30 Å; and an additional penalty of up to 36.0 Rosetta units if the inter-atom distance exceeds 30 Å. A “weak” restraint would be:

~~~
AtomPair O2′ 2A C4′ 38A FADE 0 30 15 −0.80 0.80
AtomPair O2′ 2A C4′ 38A FADE −99 60 30 −7.20 7.20
~~~

that is, one-fifth the strength of the “strong” restraint.

We show the results of the FARFAR2 ROSIE server simulations conducted with and without MOHCA-seq constraints on a *Vibrio cholerae* c-di-GMP riboswitch (PDB code: 3IRW) [29]. Unlike the original protocol using FARFAR [28], FARFAR2 does not require pre-generation of helix ensembles, manual selection of models to minimize, or manual selection of a fraction of models to cluster. The constrained simulations approach much closer to the crystal conformation: its second-lowest-energy cluster is at 5.52 Å RMSD, while the closest cluster among the top ten for the unconstrained simulation is 8.91 Å RMSD (Figure 4A). This advantage is far from chance; more than half of the models produced in the constrained simulation have RMSD less than 8.0 Å, versus less than one percent for the constrained simulation (Figure 4B). FASTA, secondary structure, and constraint files necessary to reproduce this simulation are included as an Appendix.

**Figure 4.**
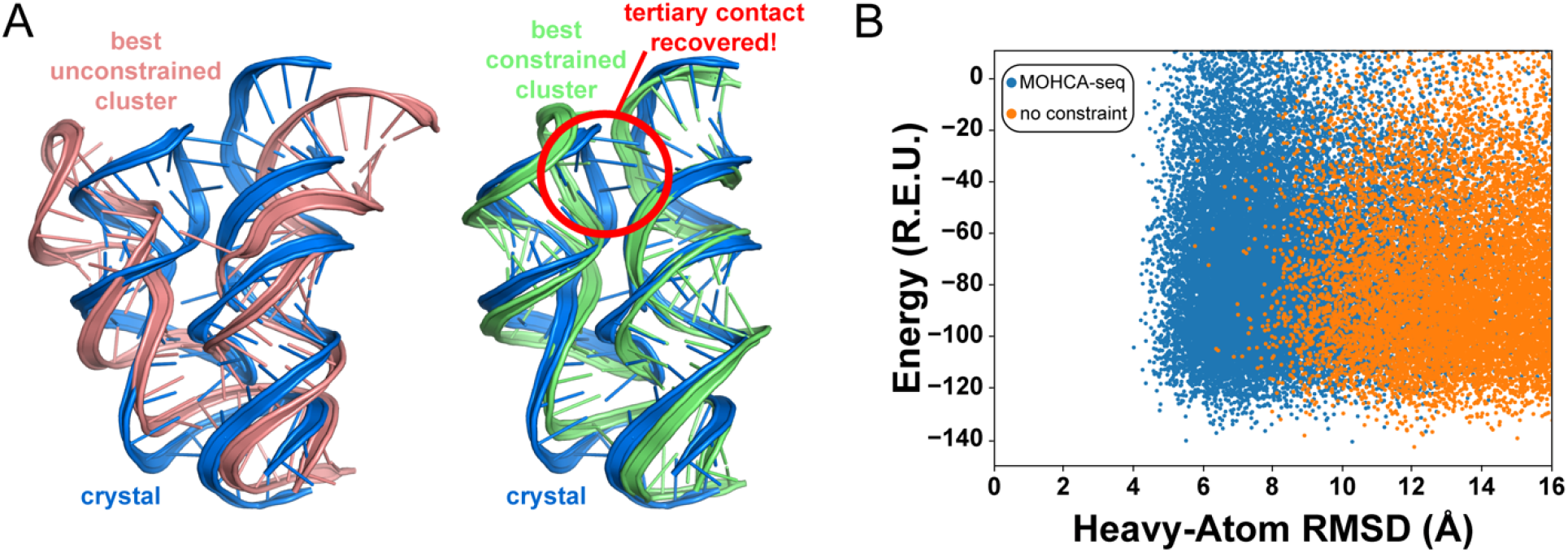
A comparison of FARFAR2 simulation results run with and without MOHCA-seq constraints on a *V. cholerae* c-di-GMP riboswitch. (A) The simulation run without the benefit of MOHCA-seq constraints is unable to recover the key tertiary contact that defines the global fold of this riboswitch (left), while a simulation with MOHCA-seq constraints finds that tertiary contact naturally (right). (B) Looking at the ensemble of generated models as a whole, a large proportion of the models from the constrained simulation are closer to the experimental model than even the best models generated in the unconstrained simulation.

#### 2.3.2 *Chemical-shift-guided FARFAR2* (CS-Rosetta-RNA)

^1^H chemical shifts alone can provide powerful information for constraining RNA folds, in many cases enabling atomic accuracy without requiring additional NMR experiments [26]. Due to improvements in the FARFAR2 protocol at baseline, the difference in performance on benchmark cases from the original CS-Rosetta-RNA study [26] is not as stark as for the earlier sampling protocol and scoring function (FARFAR rather than FARFAR2). Nonetheless, improvements from ^1^H chemical shifts remain apparent, as illustrated by the tandem GA mismatch case 1MIS [30]. A comparison of two 500-structure ensembles show that modeling guided by chemical shifts generate a much harsher score penalty for models > 2.0 Å of an experimental structure (an energy gap of 16.0 rather than 9.8 Rosetta energy units) (Figure 5).

**Figure 5.**
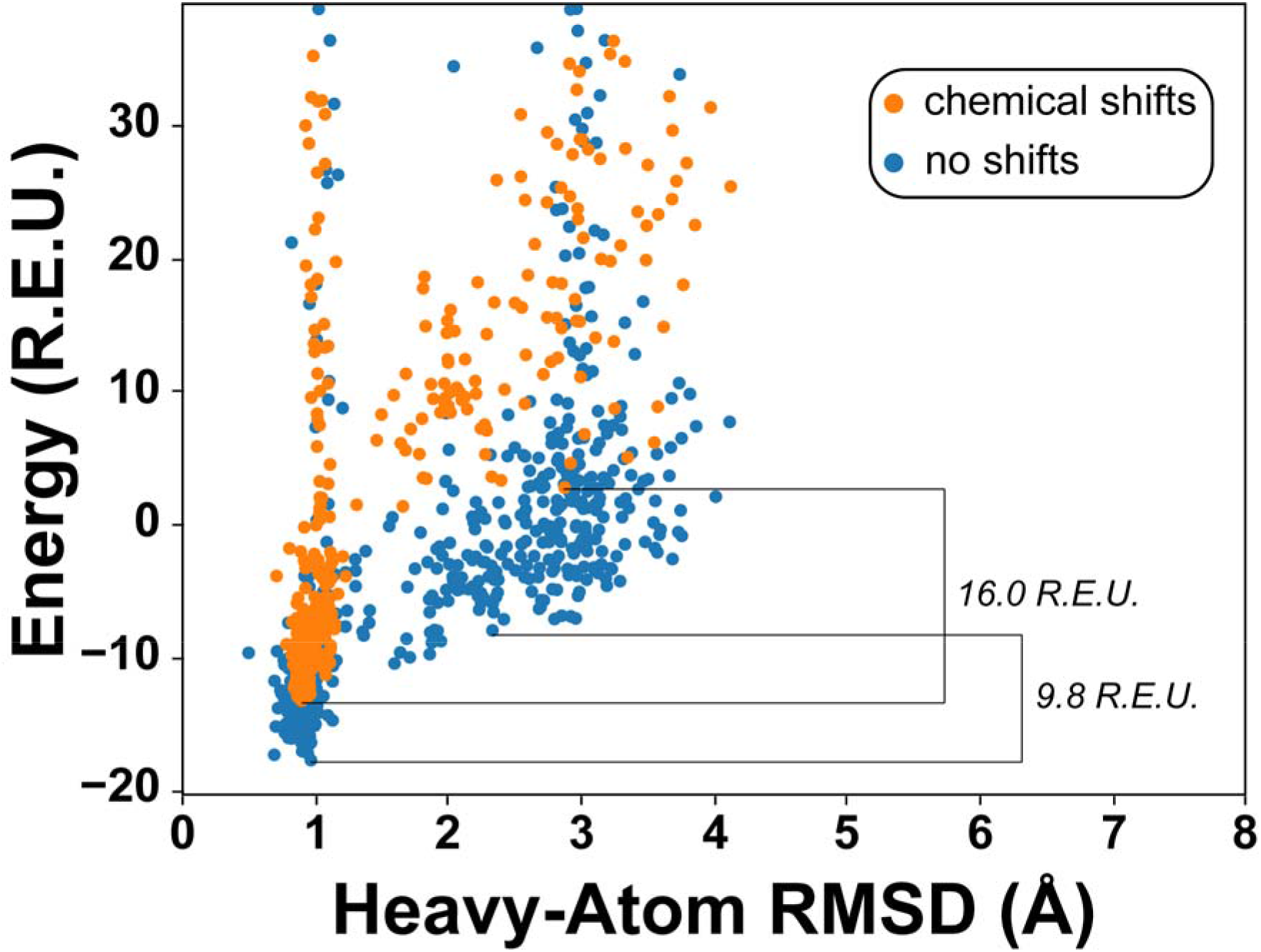
A comparison of FARFAR2 simulations run with and without chemical shifts on a duplex RNA containing tandem GA pairs. FARFAR2 can find the correct structure even without chemical shift data, but the energy function favors correct conformations by a larger margin. We show the energy gap differentiating the lowest energy correct structure (at most 1.0 Å RMSD) versus the lowest energy incorrect structure (at least 2.0 Å RMSD).

## Conclusions

The ROSIE server for FARFAR2 provides a web interface to Rosetta’s application for *de novo* modeling of complex RNA folds, targeted to users at diverse levels of expertise. Important stages of modeling, such as reducing a modeled ensemble to a handful of representative structures, are automated, enabling the user to come away with both full modeling output but also the most important models and their expected accuracy. Although the computational demands of RNA molecules remain high and atomic accuracy for intricate interactions remains difficult to achieve, the ROSIE interface ensures that non-expert can access Rosetta RNA modeling code in a state that keeps pace with ongoing developments.

## Acknowledgements

We thank Sergey Lyskov for his maintenance of the ROSIE webserver platform, lab members Rachael Kretsch, Ramya Rangan and Ivan Zheludev for server testing, and Rachael Kretsch for reading the manuscript. We acknowledge funding from the National Institutes of Health (R35 GM 122579 and R21 CA 219847)

## Appendix

All inputs to reproduce the modeling described here, as well as sample output data, are released freely in the Appendix as well as at https://github.com/DasLab/FARFAR2_modeling_examples.

## Notes

https://github.com/DasLab/FARFAR2_modeling_examples

### Competing Interest Statement

The authors have declared no competing interest.

## References

1. Cech TR, Steitz JA (2014) The noncoding RNA revolution – Trashing old rules to forge new ones. Cell 157:77–94

2. Das R, Karanicolas J, Baker D (2010) Atomic accuracy in predicting and designing noncanonical RNA structure. Nat Methods 7:291–294. https://doi.org/10.1038/nmeth.1433

3. Ditzler MA, Otyepka M, Šponer J, Walter NG (2010) Molecular dynamics and quantum mechanics of RNA: Conformational and chemical change we can believe in. Acc Chem Res. https://doi.org/10.1021/ar900093g

4. Laing C, Schlick T (2011) Computational approaches to RNA structure prediction, analysis, and design. Curr. Opin. Struct. Biol. 21:306–318

5. Cao S, Chen SJ (2005) Predicting RNA folding thermodynamics with a reduced chain representation model. RNA. https://doi.org/10.1261/rna.2109105

6. Lyskov S, Chou FC, Conchúir SÓ, et al (2013) Serverification of Molecular Modeling Applications: The Rosetta Online Server That Includes Everyone (ROSIE). PLoS One. https://doi.org/10.1371/journal.pone.0063906

7. Moretti R, Lyskov S, Das R, et al (2018) Web-accessible molecular modeling with Rosetta: The Rosetta Online Server that Includes Everyone (ROSIE). Protein Sci. https://doi.org/10.1002/pro.3313

8. Leman JK, Weitzner BD, Lewis SM, et al (2020) Macromolecular modeling and design in Rosetta: recent methods and frameworks. Nat. Methods

9. Watkins AM, Rangan R, Das R (2020) FARFAR2: Improved De Novo Rosetta Prediction of Complex Global RNA Folds. Structure. https://doi.org/10.1016/j.str.2020.05.011

10. Magnus M, Boniecki MJ, Dawson W, Bujnicki JM (2016) SimRNAweb: a web server for RNA 3D structure modeling with optional restraints. Nucleic Acids Res. https://doi.org/10.1093/nar/gkw279

11. Biesiada M, Pachulska-Wieczorek K, Adamiak RW, Purzycka KJ (2016) RNAComposer and RNA 3D structure prediction for nanotechnology. Methods. https://doi.org/10.1016/j.ymeth.2016.03.010

12. Krokhotin A, Houlihan K, Dokholyan N V. (2015) iFoldRNA v2: Folding RNA with constraints. Bioinformatics 31:2891–2893. https://doi.org/10.1093/bioinformatics/btv221

13. Parisien M, Major F (2008) The MC-Fold and MC-Sym pipeline infers RNA structure from sequence data. Nature 452:51–55. https://doi.org/10.1038/nature06684

14. Egli M, Minasov G, Su L, Rich A (2002) Metal ions and flexibility in a viral RNA pseudoknot at atomic resolution. Proc Natl Acad Sci U S A 99:4302–4307. https://doi.org/10.1073/pnas.062055599

15. Zheng L, Mairhofer E, Teplova M, et al (2017) Structure-based insights into self-cleavage by a four-way junctional twister-sister ribozyme. Nat Commun. https://doi.org/10.1038/s41467-017-01276-y

16. Leontis NB, Westhof E (2001) Geometric nomenclature and classification of RNA base pairs. RNA. https://doi.org/10.1017/S1355838201002515

17. Huang L, Lilley DMJ (2016) The Kink Turn, a Key Architectural Element in RNA Structure. J. Mol. Biol.

18. Abramovitz DL, Pyle AM (1997) Remarkable morphologlical variability of a common RNA folding motif: The GNRA tetraloop-receptor interaction. J Mol Biol. https://doi.org/10.1006/jmbi.1996.0810

19. Geary C, Baudrey S, Jaeger L (2008) Comprehensive features of natural and in vitro selected GNRA tetraloop-binding receptors. Nucleic Acids Res. https://doi.org/10.1093/nar/gkm1048

20. Fiore JL, Nesbitt DJ (2013) An RNA folding motif: GNRA tetraloop-receptor interactions. Q Rev Biophys. https://doi.org/10.1017/S0033583513000048

21. Cheng CY, Chou FC, Kladwang W, et al (2015) Consistent global structures of complex RNA states through multidimensional chemical mapping. Elife. https://doi.org/10.7554/eLife.07600

22. Smith KD, Lipchock S V., Ames TD, et al (2009) Structural basis of ligand binding by a c-di-GMP riboswitch. Nat Struct Mol Biol. https://doi.org/10.1038/nsmb.1702

23. Watkins AM, Rangan R, Das R (2019) Using Rosetta for RNA homology modeling. In: Methods in Enzymology

24. Bisaria N, Greenfeld M, Limouse C, et al (2016) Kinetic and thermodynamic framework for P4-P6 RNA reveals tertiary motif modularity and modulation of the folding preferred pathway. Proc Natl Acad Sci U S A. https://doi.org/10.1073/pnas.1525082113

25. Liu Y, Wilson TJ, Lilley DMJ (2017) The structure of a nucleolytic ribozyme that employs a catalytic metal ion. Nat Chem Biol. https://doi.org/10.1038/nchembio.2333

26. Daldrop P, Reyes FE, Robinson DA, et al (2011) Novel ligands for a purine riboswitch discovered by RNA-ligand docking. Chem Biol. https://doi.org/10.1016/j.chembiol.2010.12.020

27. Watkins AM, Geniesse C, Kladwang W, et al (2018) Blind prediction of noncanonical RNA structure at atomic accuracy. Sci Adv 4:eaar5316. https://doi.org/10.1126/sciadv.aar5316

28. Sripakdeevong P, Cevec M, Chang AT, et al (2014) Structure determination of noncanonical RNA motifs guided by 1 H NMR chemical shifts. Nat Methods. https://doi.org/10.1038/nmeth.2876

29. Wu M, Turner DH (1996) Solution structure of (rGCGGACGC)2 by twodimensional NMR and the iterative relaxation matrix approach. Biochemistry. https://doi.org/10.1021/bi960133q

